# Immunogenicity and Protective Efficacy of *Toxoplasma gondii* SRS67 and SRS20A Proteins in mice

**DOI:** 10.1101/2025.11.17.688744

**Authors:** Chen Xiaoxiao, YI Tingting, MA Wenyu, Liang Xuefei, Niu Shuaijing, Li Hongwei, Du Mengze, Zhou Shuanghai, LI Qiuming, Yin Deqi

**Affiliations:** Beijing Key Laboratory of Traditional Chinese Veterinary Medicine, Beijing University of Agriculture, Changping, Beijing, PR China; Coccidia Laboratory, College of Animal Science and Technology/College of Veterinary Medicine, Beijing University of Agriculture, Beijing, PR China; Police dog Technology Teaching and Research Section, Department of Criminal Science and Technology, Beijing Police College, Beijing, PR China

**Keywords:** *Toxoplasma gondii*, SRS67, SRS20A, immunogenicity, immune protection

## Abstract

*Toxoplasma gondii*, a medically significant zoonotic protozoan, relies on surface SAG-family antigens for host adhesion and immune evasion. SRS67 and SRS20A, two such antigens, are hypothesized to be highly immunogenic. Here, we comprehensively evaluated their immunological properties and protective efficacy in mice. Recombinant SRS67 and SRS20A, produced via prokaryotic expression, induced high-titer, specific polyclonal antisera in BALB/c mice and rabbits. Western blotting confirmed these antisera recognized native RH strain tachyzoite antigens, validating strong immunogenicity. qPCR showed SRS67 transcription was higher in PRU (chronic) than RH (acute) strains, while SRS20A expression was comparable. Immunofluorescence, using GAP45 as a membrane marker, localized both proteins as puncta around RH tachyzoite membranes. In vitro phenotypic assays indicated that the corresponding polyclonal antibodies significantly inhibited the invasion of host cells by *T. gondii* and subsequent intracellular proliferation. Challenge studies demonstrated protective efficacy: in acute RH infection, immunized mice (single or combined antigens) showed 60% higher survival, with bivalent groups delaying mortality by 1-2 days. In chronic PRU infection, all immunized groups had >85% fewer brain cysts, and bivalents better attenuated neuropathology (e.g., meningeal thickening). Mechanistically, SRS67 upregulated Th1 cytokines (TNF-α, IL-18) in acute infection, while both proteins downregulated IFN-γ to limit immune-mediated damage; GSH upregulation indicated antioxidative involvement. In chronic infection, co-immunization synergistically enhanced SOD activity to counter oxidative stress, with IFN-γ downregulation maintaining immune homeostasis. These findings support SRS67 and SRS20A as promising *T. gondii* vaccine candidates, acting via coordinated Th1 immunity and antioxidative pathways.

## Introduction

*Toxoplasma gondii* is an obligate intracellular apicomplexan parasite[1]. Its heteroxenous life cycle alternates between felids—its definitive hosts—and virtually all warm-blooded vertebrates, including humans, which serve as intermediate hosts[2,3]. Transmission routes of toxoplasmosis are diverse and include congenital (vertical) transmission; food- or water-borne acquisition of oocysts or tissue cysts, percutaneous or mucosal inoculation, and iatrogenic infection via blood transfusion or organ transplantation[4]. Pathogenicity is largely dictated by the host’s immune status: whereas immunocompetent individuals usually remain asymptomatic, immunocompromised patients may develop life-threatening complications such as toxoplasmic encephalitis or chorioretinitis[5,6]. In livestock, *T. gondii* infection is a major cause of abortion and stillbirths, resulting in substantial economic losses[7]. Current therapeutic regimens rely primarily on pyrimethamine combined with sulfonamides; however, these agents are associated with considerable toxic side effects[8]. Consequently, there is an urgent need to develop novel antitoxoplasmic drugs and effective vaccines.

The surface antigen 1-related sequence (SRS) proteins of *Toxoplasma gondii* are a family of surface-anchored membrane proteins that share high structural homology with the archetypal surface antigen SAG1. They govern the entire cascade of parasite recognition, adhesion, and host-cell invasion, rendering them ideal targets for vaccine development[9]. Recent functional studies have begun to dissect the roles of individual SRS members: deletion of SRS19d markedly reduces plaque size, tachyzoite numbers, virulence, and cyst burden[10], whereas loss of P18 (SRS35/SAG4) attenuates virulence yet paradoxically enhances cyst formation and stimulates host immunity[11]. The flagship antigen SAG1 has provided a proof of concept: its recombinant protein elicits potent Th1-biased responses and significantly prolongs survival in murine models[12], and its DNA vaccine markedly diminishes brain cyst loads[13]. Likewise, SRS52 exhibits robust immunogenicity and markedly improves the survival rate of infected mice[14]; comprehensive bioinformatic analyses of SRS40E[15] and SRS47D[16], alongside the Vax452 vaccine platform based on SRS proteins, further highlight the potential of protein family in *anti-Toxoplasma* vaccination[17]. Nevertheless, single-antigen formulations rarely confer complete protection. Encouragingly, multi-antigen strategies have demonstrated synergistic efficacy: a triple vaccine composed of TgMIC1-4-6 elevated mouse survival to 80% and reduced parasite burden by two orders of magnitude[18]. Collectively, these findings strongly suggest that rational combination of multiple SRS antigens may overcome current limitations in vaccine efficacy and open new avenues for next-generation vaccines against toxoplasmosis.

In the present study, we evaluated the protective capacity of recombinant SRS67 and SRS20A—administered individually or as a bivalent formulation—against acute (RH strain) and chronic (PRU strain) *T. gondii* infection in BALB/c mice.

## Materials and Methods

### Cells and parasites

Vero cells were maintained in 25 cm² culture flasks with DMEM (Servicebio, China) containing 100 U/mL penicillin, 100 µg/mL streptomycin (Leagene, China), and 10% heat-inactivated fetal bovine serum (FBS; BBI, China) at 37°C with 5% CO₂. *Toxoplasma gondii* tachyzoites of the RH strain were propagated in Vero cells under identical incubation conditions, using the same medium with FBS reduced to 2%.

### Molecular cloning and expression of recombinant SRS67 and SRS20A proteins

Total RNA was extracted from *T. gondii* RH tachyzoites with TRIzol (Beyotime, Shanghai, China) and reverse-transcribed into cDNA. Gene-specific primers containing restriction sites were designed on the basis of ToxoDB sequences and synthesized by Sangon Biotech (Shanghai): (*SRS67*, F: 5’-cgGGATCCatgaagagtgttctccacagcg-3’, R: 5’-ccgCTCGAGtcaaagtgtttgcgacacgctt-3’; *SRS20A*, F: 5’-cGAGCTCatggcggatccagaaaagtgtg-3’, R: 5’-cccAAGCTTtcacaacattttcatcaacacgacag-3’). PCR products were ligated into pUCm-T vectors (Sangon Biotech, Shanghai, China) and sequence-verified. Correct inserts were released by BamHI/XhoI (for *SRS67*; Beyotime, Shanghai, China) or SacI/HindIII (for *SRS20A*; Beyotime, Shanghai, China) digestion and subcloned into pET-32a, followed by re-sequencing for confirmation. The detailed experimental protocols were adopted from those previously established by our laboratory[19].

### Preparation of recombinant SRS67 and SRS20A proteins

Recombinant plasmids pET32a-SRS67 and pET32a-SRS20A were individually transformed into *E. coli* BL21 (DE3; Sangon Biotech, Shanghai, China) competent cells. Positive clones were cultured in LB containing 100 µg/mL ampicillin at 37°C until OD₆₀₀ ≈ 0.6, then induced with 0.1 or 0.5 mM IPTG for 16 h at the same temperature. Cells were harvested by centrifugation (5,000 g/min, 10 min, 4°C), resuspended in ice-cold PBS (pH 7.4) supplemented with 1 mM PMSF, and lysed by sonication on ice (3 s pulse, 5 s interval, 10 min total). After centrifugation (15,000 g/min, 30 min, 4°C), the pellets containing inclusion bodies were solubilized, and proteins were purified under denaturing conditions using Ni-NTA agarose beads (Beyotime, Shanghai, China) according to the manufacturer’s protocol. Purified proteins were refolded by stepwise dialysis against buffers containing decreasing urea concentrations (6 → 0 M) followed by decreasing arginine concentrations (0.5 → 0 M) at 4°C (Sangon Biotech, Shanghai, China). Refolded proteins were concentrated, flash-frozen in liquid nitrogen, and stored at −80°C until use.

Recombinant TgSRS67 and TgSRS20A were resolved on 10% SDS-PAGE gels and transferred (180 mA, 88 min, 4°C) onto 0.22 µm PVDF membranes (Bio-Rad, Hercules, CA, USA). Membranes were blocked with 5% (w/v) BSA (Yuanye, Shanghai, China) in TBS-T (20 mM Tris-HCl, pH 7.6, 150 mM NaCl, 0.05% Tween-20) for 1 h at room temperature, incubated overnight at 4°C with mouse anti-His monoclonal antibody (1:5,000; Solarbio, Beijing, China), washed three times (10 min each) with TBS-T, and probed with HRP-conjugated goat anti-mouse IgG (1:10,000; Beyotime, Shanghai, China) for 1 h at RT. Immunoreactive bands were visualized using BeyoECL Star substrate (Beyotime, Shanghai, China) and documented on a gel imaging system (Clinx, Shanghai, China).

### Polyclonal antisera production

Six-week-old female BALB/c mice (n = 6) and New Zealand white rabbits (n = 6) were randomly assigned to SRS67, SRS20A, or PBS control groups. Recombinant proteins were emulsified 1:1 (v/v) with either Freund’s Complete Adjuvant (FCA; Beyotime, Shanghai, China) for priming or Freund’s Incomplete Adjuvant (FIA; Beyotime, Shanghai, China) for booster immunizations. Mice received 50 µg, and rabbits 200 µg, of protein per dose. Immunizations were performed subcutaneously on days 0, 14, 28, and 42. Seven days after the final boost, sera were collected and screened by indirect ELISA for endpoint titers of ≥ 1:32 000. Qualified animals were terminally bled (orbital plexus for mice, cardiac puncture for rabbits) under isoflurane anesthesia. Sera were heat-inactivated (56°C, 30 min), clarified (12,000 g/min, 10 min, 4°C), aliquoted, and stored at −80°C until use. The specificity of the antisera was confirmed by Western blot against *T. gondii* RH tachyzoite lysate.

### IgG purification

The BeyoMag™ Protein A+G magnetic beads (Beyotime, Shanghai, China; were washed three times each with PBS (10 mM Na₂HPO₄, 2 mM KH₂PO₄, 137 mM NaCl, 2.7 mM KCl, pH 7.4) and once with 1 mL 50 mM sodium phosphate (pH 7.0). Serum and sodium phosphate were added and incubated at 4°C for 1 h. After two washes, IgG was eluted at 4°C with citric acid three times. The supernatants were immediately neutralized to pH 7.0 with Tris-HCl (pH 9.0). Combined eluates were dialyzed against PBS for 2 h, quantified, stored at –80°C, and analyzed by SDS-PAGE.

### Immunofluorescence assay

Vero cells were seeded on coverslips for 24 h, then infected with *T. gondii* RH tachyzoites. After 2 h, unattached parasites were removed by PBS washing, and cultures were continued until parasitophorous vacuoles formed. Cells were fixed with 4% paraformaldehyde (Labgic, Beijing, China) for 30 min, permeabilized with 0.15% Triton X-100 (Solarbio, Beijing, China) for 15 min, and blocked with 5% BSA for 1 h. Coverslips were incubated overnight at 4°C with purified anti-SRS67 or anti-SRS20A polyclonal antibody (Solarbio, Beijing, China), washed three times with PBST, and then treated with fluorescent secondary antibody for 1 h at 37°C in the dark. Nuclei were stained with DAPI (Beyotime, Shanghai, China) for 5 min, mounted with antifade medium, and protein localization was examined using fluorescence microscopy (Olympus, Tokyo, Japan).

### Quantitative transcriptional analysis of SRS67 and SRS20A genes

The sequences of *GAPDH*, *SRS67* and *SRS20A* were downloaded and sent to Sangon Biotech (Shanghai) for qPCR primer synthesis: *GAPDH*, F: 5’-TGGTGTTCCGTGCTGCGATGGAAC-3’, R: 5’-GAGCTTGCCGTCCTTGTGGCTGAC-3’; *SRS67*, F: 5’-GGGCGACGGCGACAATTCC-3’, R: 5’-GCATTCCCTCCAGTCCAGAACAG-3’; *SRS20A*, F: 5’-AACGCACAGATGAAAGGGGAAGC-3’, R: 5’-GTCCATGCTCCGCTCGTGATTC-3’. Using cDNA from *T. gondii* RH and PRU strains, qPCR was performed in 50 μL containing 25 μL 2× FastSYBR Mixture (Cwbio, Jiangsu, China), 1 μL each primer, 2 μL cDNA, and ddH₂O to 50 μL. Cycling conditions: 95°C for 20 s; 35 cycles of 95°C for 3 s, 60°C for 30 s; followed by melt-curve 95°C for 15 s, 60°C for 1 min, 95°C for 15 s, 60°C for 15 s.

### Plaque assay

Vero cells were evenly seeded into 12-well plates and incubated overnight in a cell culture incubator. 1 × 10³ tachyzoites were pre-incubated with anti-SRS67 or anti-SRS20A polyclonal antibodies at 0, 10, 50, or 100 µg/mL, then used to infect the Vero cell monolayers. After approximately 7 days, cells were stained with crystal violet (Solarbio, Beijing, China), and the number and area of plaques were recorded. Plaque areas were quantified with ImageJ, while plaque counts were statistically analyzed using GraphPad Prism 10.

### Invasion assay

Freshly egressed *T. gondii* RH (1 × 10⁶ tachyzoites) were pre-incubated with serial dilutions of purified anti-SRS67 or anti-SRS20A IgG (0–100 μg mL⁻¹) in DMEM containing 1% FBS for 30 min at 37°C. Vero cell monolayers (2 × 10⁵ cells per well, 24-well plates) were inoculated with the antibody-parasite mixtures and centrifuged (300 g/min, 5 min, RT) to synchronize invasion. After 3 h at 37°C, non-invaded parasites were removed by washing three times with PBS, and cells were fixed with 4% paraformaldehyde (10 min). Fixed monolayers were stained with 10% Giemsa (Beyotime, Shanghai, China) for 20 min, rinsed with distilled water, and air-dried. Twenty randomly selected fields per well were imaged at 40× magnification using a microscope. The invasion rate (%) was calculated as (number of parasitophorous vacuoles/total host nuclei) × 100 and averaged across three independent experiments, each performed in triplicate.

### Proliferation assay

The procedure for the proliferation assay was similar to that of the invasion assay. After the 3 h invasion period and subsequent washing to remove extracellular parasites, the infected monolayers were further incubated in fresh medium for an additional 24 h to allow for intracellular replication. The cells were then fixed, and the number of parasites per vacuole was determined by examining at least 100 parasitophorous vacuoles per well under a 40× objective lens.

### Egress assay

The procedure for the egress assay was similar to that of the invasion assay. After the 3 h invasion period, non-invaded parasites were removed with PBS. Cultures were continued until parasitophorous vacuoles became prominent, then washed twice more with PBS. Serum IgG was added at 0, 10, 50, or 100 μg/mL (triplicate wells per dose). Incubation proceeded until obvious parasite egress was observed. The supernatant was collected, centrifuged at 3,000 rpm for 5 min, and the pellet was resuspended in PBS for enumeration of egressed parasites.

### Immunoprotection experiment

*T. gondii* RH and PRU strains were each allocated to five groups (10 mice per group): RH/PRU infection control, SRS67 single-dose (50 μg), SRS20A single-dose (50 μg), SRS67+SRS20A combined, and PBS blank. Mice were immunized according to the established protocol using Freund’s adjuvant. After immunization, all groups except the PBS blank control were challenged with 1 × 10² RH tachyzoites (intraperitoneal injection) or 15 PRU cysts (oral gavage). Survival was monitored for >30 days post-challenge (survival rate = number of survivors/10 × 100%). Upon completion, PRU-strain mice were euthanized; brain cysts were counted in three mice per group, and the remaining brains were fixed in 4% paraformaldehyde for 24 h and sent to Wuhan Servicebio for histopathological analysis.

### Determination of Oxidative Factors and Immune Factors Levels

The grouping, immunization, and challenge procedures for this experiment were consistent with those described in section 2.12 (with the exception that the T. gondii RH strain groups lacked the SRS67+SRS20A combined immunization group). On day 7 post-challenge, 3 mice were randomly selected from each group of both RH strain-challenged and PRU strain-challenged mice, respectively. These mice were euthanized, and their spleen tissues were promptly harvested. Total RNA was extracted from the spleen tissues. Subsequent qPCR analysis was performed to determine the expression levels of cytokines and antioxidant stress factors. The specific primers for the target cytokines and antioxidant stress factors involved in this detection were synthesized by Sangon Biotech (Shanghai, China): GAPDH, F: 5’-ATCAAGAAGGTGGTGAAGCAGG-3’, R: 5’-GAAGAGTGGGAGTTGCTGTTGAA G-3’; SOD, F: 5’-CCAGACCTGCCTTACGACTATG-3’, R: 5’-CTCGGTGGCGTTGAGATTGT-3’; GSH, F: 5’-CAGTCCACCGTGTATGCCTTC-3’, R: 5’-CTCCTGGTGTCCGAACTGAT-3’; CAT, F: 5’-GGAGGCGGGAACCCAATAG-3’, R: 5’-GTGTGCCATCTCGTCAGTGAA-3’; IFN-γ, F: 5’-ACAATCAGGCCATCAGCAACAAC-3’, R: 5’-ATTGAATGCTTGGCGCTGGAC-3’; TNF-α, F: 5’-CCTGTAGCCCACGTCGTAG-3’, R: 5’-GGGAGTAGACAAGGTACAACCC-3’; IL-18, F: 5’-GACTCTTGCGTCAACTTCAAGG-3’, R: 5’-CAGGCTGTCTTTTGTCAACGA-3’; IL-10, F: 5’-CGGGAAGACAATAACTGCACCC-3’, R: 5’-CGGTTAGCAGTATGTTGTCCAGC-3’; IL-4, F: 5’-GGTCTCAACCCCCAGCTAGT-3’, R: 5’-GCCGATGATCTCTCTCAAGTGAT-3’.

### Evolutionary tree analysis

SRS67 and SRS20A nucleotide and protein sequences were retrieved from ToxoDB (https://toxodb.org/toxo/app) to construct a phylogenetic tree. Homologs with the highest BLAST scores were selected (for SRS67, a total of 20 sequences were included, nt Score > 250, AA Score > 240; for SRS20A, a total of 17 sequences were included, nt Score > 240, AA Score > 280). Nucleotide and amino acid alignments were generated with MEGA 11, and neighbor-joining trees with pairwise genetic-distance estimates were constructed under default parameters. The evolutionary tree diagram were edited in ITOL (https://itol.embl.de). Nucleotide and protein identity matrices derived from the BLAST outputs were visualized with TBtools-II. Finally, the domains (https://www.ncbi.nlm.nih.gov/Structure/cdd/wrpsb.cgi) and tertiary structures (https://swissmodel.expasy.org/interactive) of the selected sequences mentioned above were summarized and analyzed.

### Statistical analysis

All data were analyzed and graphed using GraphPad Prism 10.3.0 (GraphPad Software, San Diego, CA, USA). Transcript levels between strains were compared with unpaired, two-tailed Student’s *t*-tests. Cellular assays were evaluated by one-way or two-way ANOVA followed by Tukey’s multiple-comparisons test as appropriate. Results are presented as mean ± SD from at least three independent experiments. Statistical significance was defined as ns, *P* > 0.05; *, *P* < 0.05; **, *P* < 0.01; and ***, *P* < 0.001.

## Results

### Generation and validation of anti-SRS67 and anti-SRS20A polyclonal antibodies

Based on our previous bioinformatic predictions, *Toxoplasma gondii* SRS67 and SRS20A proteins exhibit high hydrophilicity, favorable thermostability, and abundant B-cell and CTL epitopes. In this study, we successfully constructed pET32a-SRS67 and pET32a-SRS20A recombinant plasmids [19]. After IPTG-induced expression and affinity purification, Western blot analysis revealed specific bands at approximately 39.5 kDa for SRS67 and 51.3 kDa for SRS20A, consistent with their theoretical molecular weights (kDa), indicating that highly pure and intact recombinant proteins were obtained (Fig. 1A-D). The prediction results showed that both proteins had signal peptide regions and SAG super family domains (Fig. 1E).

**Fig. 1.**
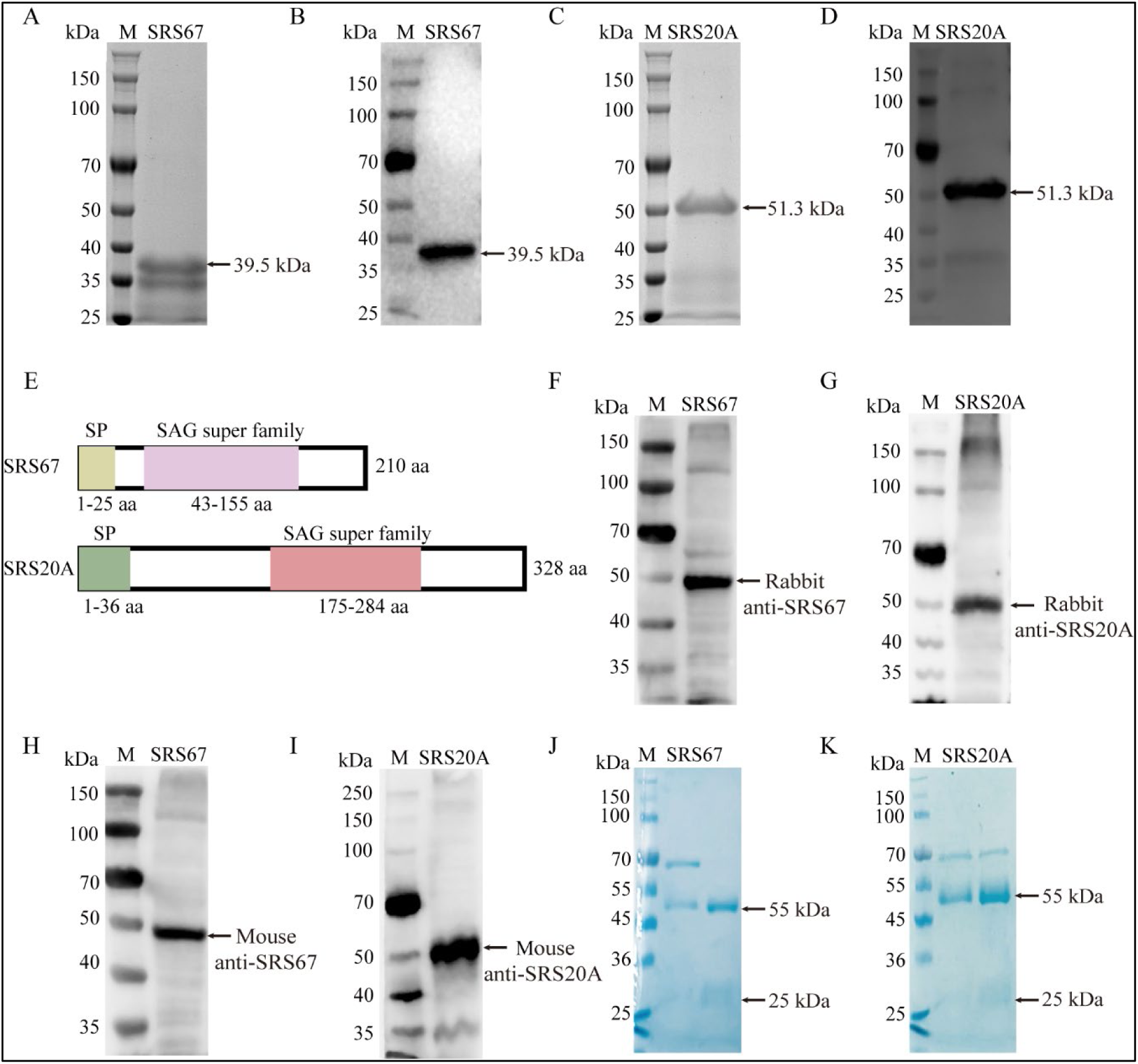
Protein structure model prediction and antibody specificity detection. **A** Protein purification result of SRS67 (SDS-PAGE). **B** Protein purification result of SRS67 (Western blot). **C** Protein purification result of SRS20A (SDS-PAGE). **D** Protein purification result of SRS20A (Western-blot). **E** Protein domain prediction of SRS67 and SRS20A. SP, signal peptide. **F** Western blot analysis of native TgSRS67 expression using rabbit polyclonal antibodies. **G** Western blot analysis of native TgSRS20A expression using rabbit polyclonal antibodies. **H** Western blotting analysis of native TgSRS67 expression using mouse polyclonal antibodies. **I** Western blotting analysis of native TgSRS20A expression using mouse polyclonal antibodies. **J** SDS-PAGE of purified serum IgG from SRS67. **K** SDS-PAGE of purified serum IgG from SRS20A.

Following immunization, sera from both BALB/c mice and New Zealand white rabbits exhibited endpoint ELISA titers ≥ 1:32,000 against SRS67 and SRS20A (Fig. S1), indicating robust humoral responses and high functional avidity. Western blot analyses demonstrated that murine and rabbit anti-SRS67/SRS20A antibodies specifically recognized native TgSRS67 and TgSRS20A in RH-strain tachyzoite lysates without detectable cross-reactivity (Fig. 1F-I).

Polyclonal IgG was subsequently purified from pooled immune rabbit sera by Protein A/G magnetic bead chromatography. BCA quantification yielded concentrations of 1.36 μg/mL (anti-SRS67) and 2.14 μg/mL (anti-SRS20A). Under reducing SDS-PAGE, each preparation displayed single, sharp bands at approximately 55 kDa (heavy chain) and 25 kDa (light chain) (Fig. 1J-K), confirming antibody integrity and >95% purity.

### Transcriptional analysis of SRS67 and SRS20A in *T. gondii* strains

Quantitative RT-PCR revealed that SRS67 transcript abundance was markedly higher in the PRU strain than in the RH strain (*P* < 0.001), whereas SRS20A expression did not differ significantly between the two strains (*P* > 0.05) (Fig. S2).

### Subcellular localization of SRS67 and SRS20A

Immunofluorescence analysis showed that, using the *T.gondii* membrane protein GAP45 as a marker, revealed that SRS67 and SRS20A proteins were mainly distributed in a punctate pattern around the tachyzoite membrane of *Toxoplasma gondii* (Fig. 2A-B).

**Fig. 2.**
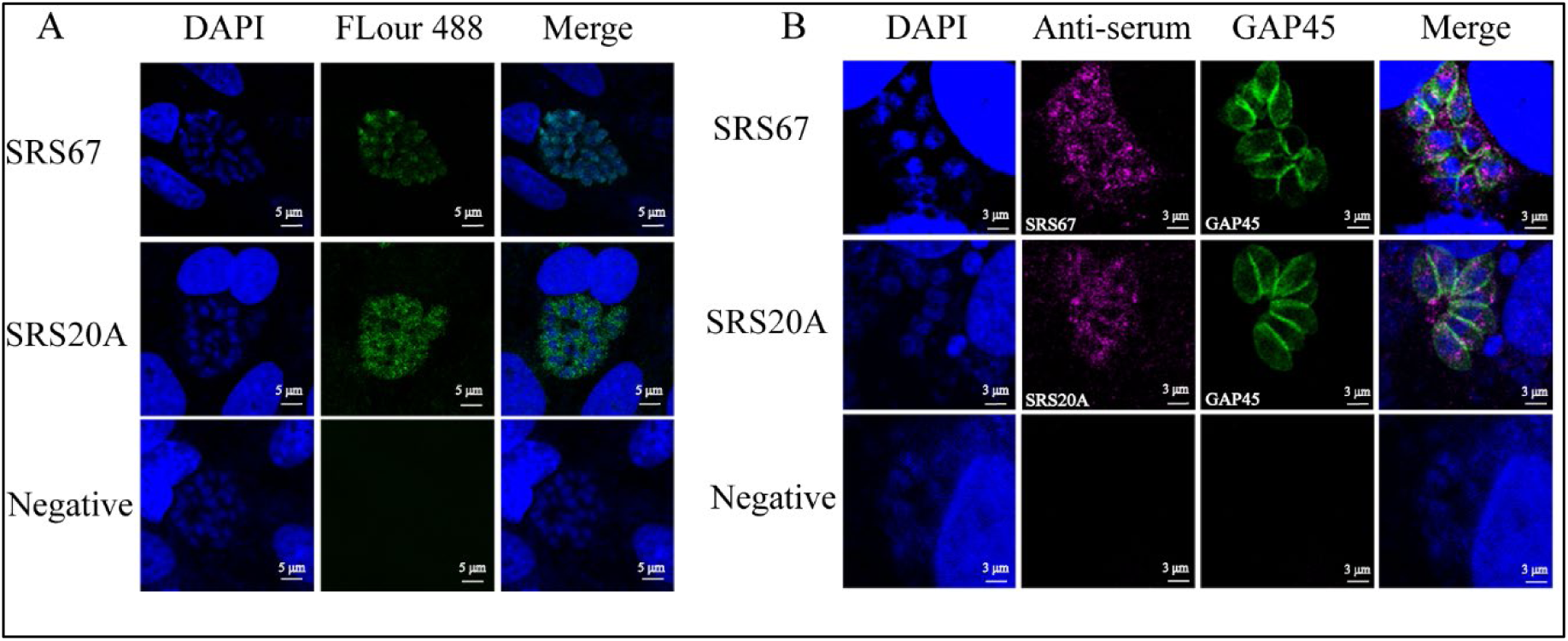
Localization of SRS67 and SRS20A proteins in *Toxoplasma gondii.* **A** Indirect immunofluorescence of SRS67 and SRS20A. **B** Co-localization of SRS67, SRS20A and GAP45.

### Anti-SRS67 and anti-SRS20A antibodies inhibit *T. gondii* invasion, replication, and egress

Cytotoxicity assays confirmed that both anti-SRS67 and anti-SRS20A IgG were well tolerated, with Vero cell viability exceeding 90% at concentrations up to 100 μg/mL (Fig. 3A-B). Plaque assays revealed a concentration-dependent reduction in both plaque number and area. Relative to untreated controls, 50 and 100 μg/mL anti-SRS67 IgG decreased plaque number by 39% and 65% (Fig. 3C-D; *P* < 0.001), respectively, while anti-SRS20A IgG produced 43% and 69% reductions (Fig. 3D-E; *P* < 0.001). Invasion assays demonstrated that both antibodies significantly impaired parasite entry in a dose-dependent manner. At 100 μg/mL, anti-SRS67 and anti-SRS20A IgG reduced invasion rates to 43% and 51% of control values, respectively (Fig. 3F-G; *P* < 0.001). The proliferation experiments showed that both anti-SRS67 and anti-SRS20A antibodies significantly inhibited the proliferation of *Toxoplasma gondii* within the host cells. In the control group (0 μg/mL), the parasitophorous vacuoles primarily contained 8 (approximately 35%) and 16 (approximately 53%) tachyzoites, while in the antibody-treated groups, they were mainly composed of 4, 8, and 16 tachyzoites. When the anti-SRS67 antibody was applied at concentrations ranging from 0 to 100 μg/mL, the proportion of parasitophorous vacuoles (PVs) containing 8 tachyzoites significantly increased from 35% to 59% (*P*<0.001), while that of PVs containing 16 tachyzoites significantly decreased from 53% to 13% (Fig. 3H; *P*<0.001). The anti-SRS20A antibody showed a similar trend: the proportion of PVs containing 8 tachyzoites increased from 35% to 57% (*P*<0.001), and that of PVs containing 16 tachyzoites decreased from 53% to 20.3% (Fig. 3I; *P*<0.001). Egress assays indicated that anti-SRS67 IgG suppressed parasite egress in a concentration-dependent fashion, with 50 and 100 μg/mL reducing egress by 51% and 81% (Fig. 3J; *P* < 0.001). In contrast, anti-SRS20A IgG required 100 μg/mL to achieve significant inhibition (Fig. 3K; *P* < 0.001).

**Fig. 3.**
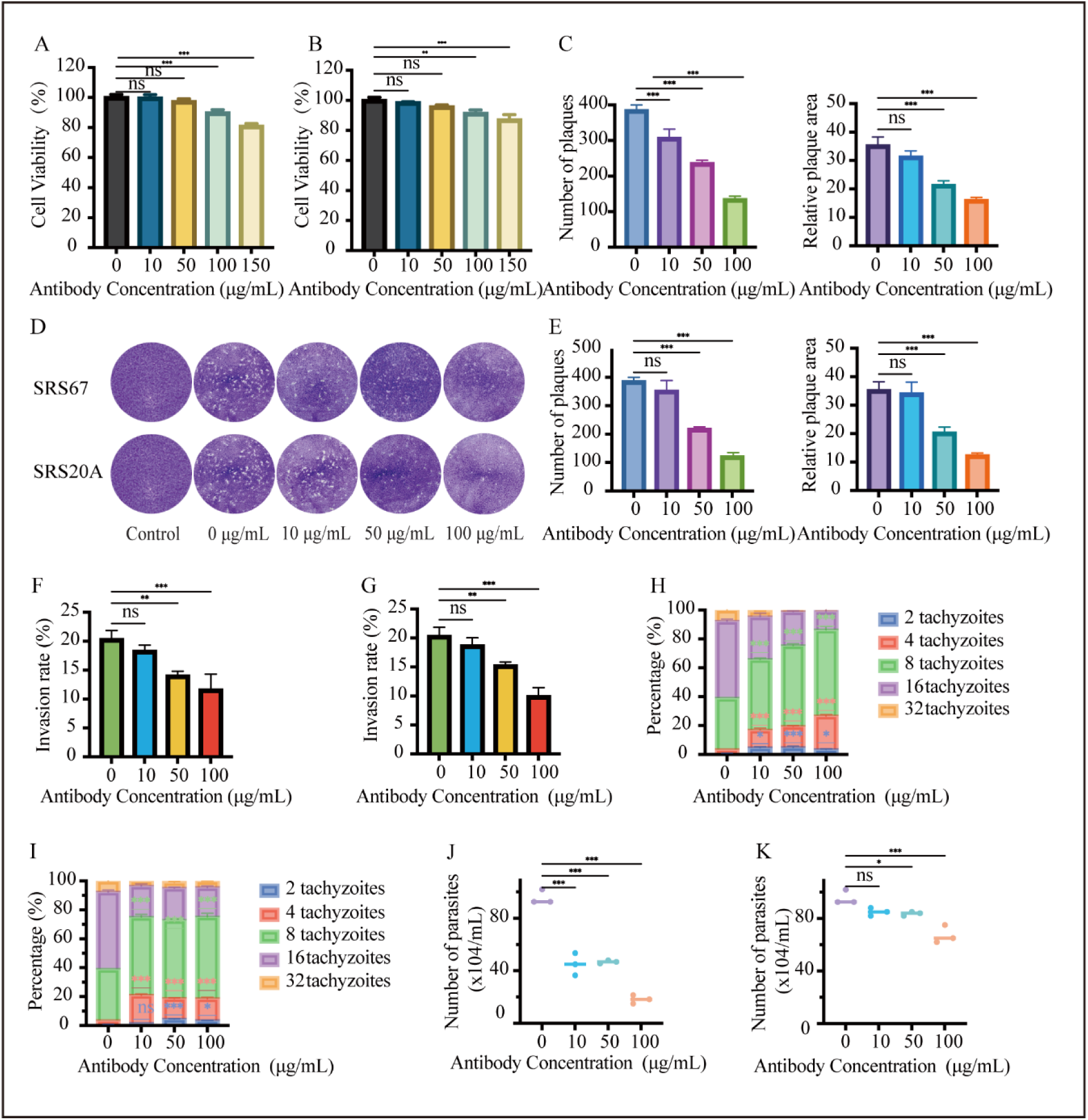
Antibody Inhibition Experiment. **A** The safe concentration of anti-SRS67 antibody - cell survival rate. **B** The safe concentration of anti-SRS20A antibody - cell survival rate. **C** The influence of SRS67 antibody on the area and quantity of plaques. **D** The effect of SRS67 and SRS20A antibodies on the formation of plaques by the parasite. **E** The influence of SRS20A antibody on the area and quantity of plaques. **F** The effect of anti-SRS67 antibody on the invasion of *T. gondii*. **G** The effect of anti-SRS20A antibody on the invasion of *T. gondii*. **H** The effect of anti-SRS67 antibody on the proliferation of *T. gondii*. **I** The effect of anti-SRS20A antibody on the proliferation of *T. gondii*. **J** The effect of anti-SRS67 antibody on the escape of *T. gondii*. **K** The effect of anti-SRS20A antibody on the escape of *T. gondii*.

### Immunoprotection against *T. gondii* infection

Following the immunization protocol, mice were challenged with *Toxoplasma gondii* and survival was monitored (Fig. 4A). In the acute-infection model (RH strain), all untreated control mice succumbed within 10 days post-challenge, whereas mice immunized with SRS67, SRS20A, or the combined formulation exhibited a 60% survival rate. Notably, the bivalent vaccine group showed a 1-2 day delay in mortality compared with the monovalent vaccine groups (Fig. 4B). In the chronic-infection model (PRU strain), only 80% of untreated controls survived, whereas 100% of the SRS67-, SRS20A- and combined-immunized mice remained alive (Fig. 4C). These results demonstrate that SRS67 and SRS20A significantly enhance resistance to *T. gondii*, although the bivalent vaccine did not improve final survival in acute infection.

**Fig. 4.**
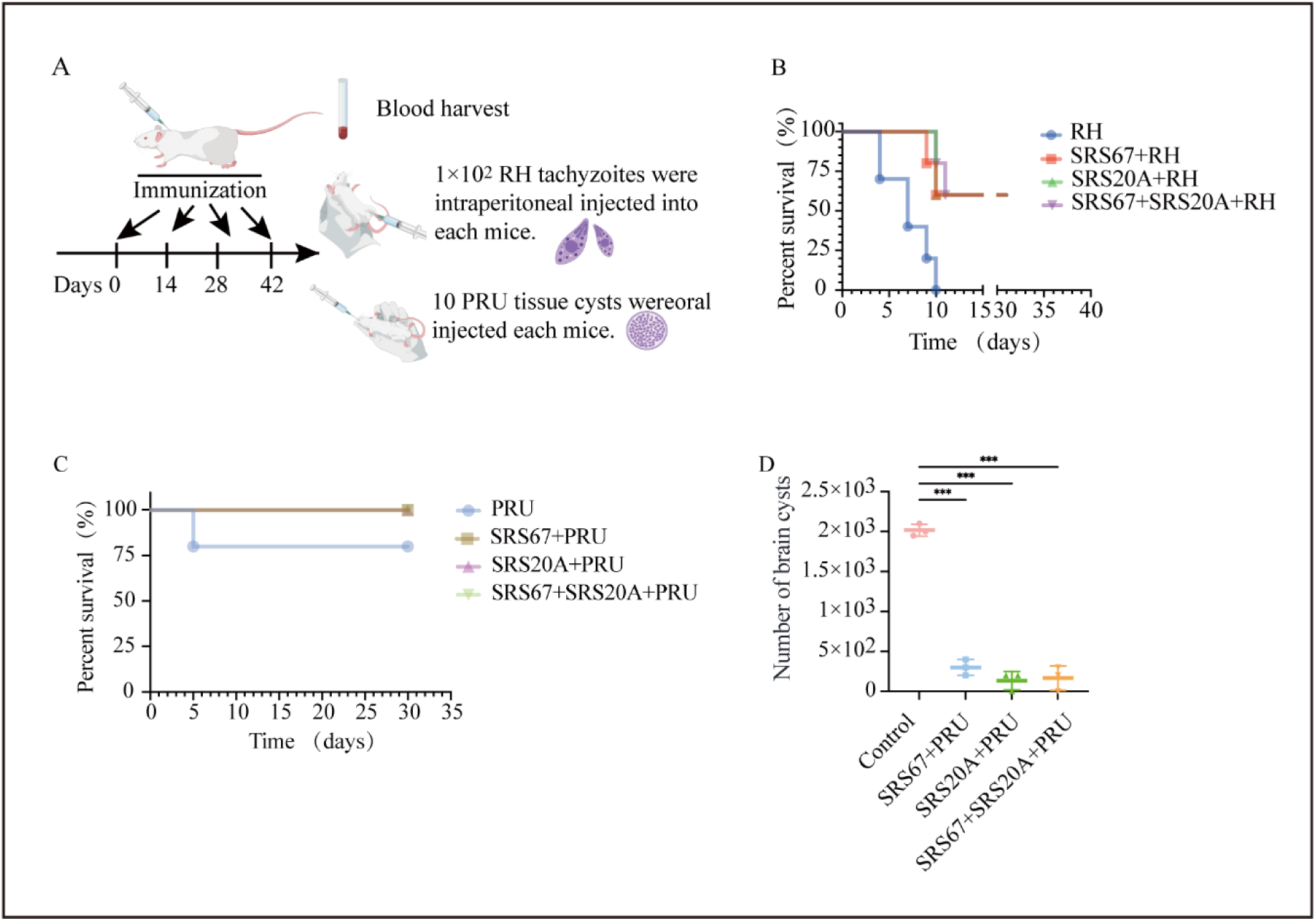
The protective effects of SRS67 and SRS20A proteins on mice infected with *Toxoplasma gondii*. **A** Schematic diagram of mouse protein immunization and *Toxoplasma gondii* infection. **B** Survival rate of mice infected with RH strain after immunization with protein. **C** Survival rate of mice infected with PRU strain after immunization with protein. **D** Changes in the number of brain cysts after protein immunization.

Additionally, brain cyst burden and histopathological changes were assessed in PRU-strain-infected mice. As shown in Fig. 4D, compared with the PRU-infection group, the number of brain cysts was reduced by 85.1% in the SRS67+PRU group, by 93.3% in the SRS20A+PRU group, and by 91.7% in the SRS67+SRS20A+PRU group (*P* < 0.001). Histopathological examination of brain sections (Fig. 5) showed severe lesions in non-immunized animals, including perivascular cuffing (black arrows), meningeal thickening (blue arrows), neuronal necrosis with neuronophagia (green arrows), and submeningeal inflammatory infiltrates (yellow arrows). In contrast, vaccinated mice displayed only mild inflammatory changes, and the bivalent vaccine group exhibited slightly attenuated neuropathology relative to the monovalent groups, indicative of an additive, but not synergistic, protective effect.

**Fig. 5.**
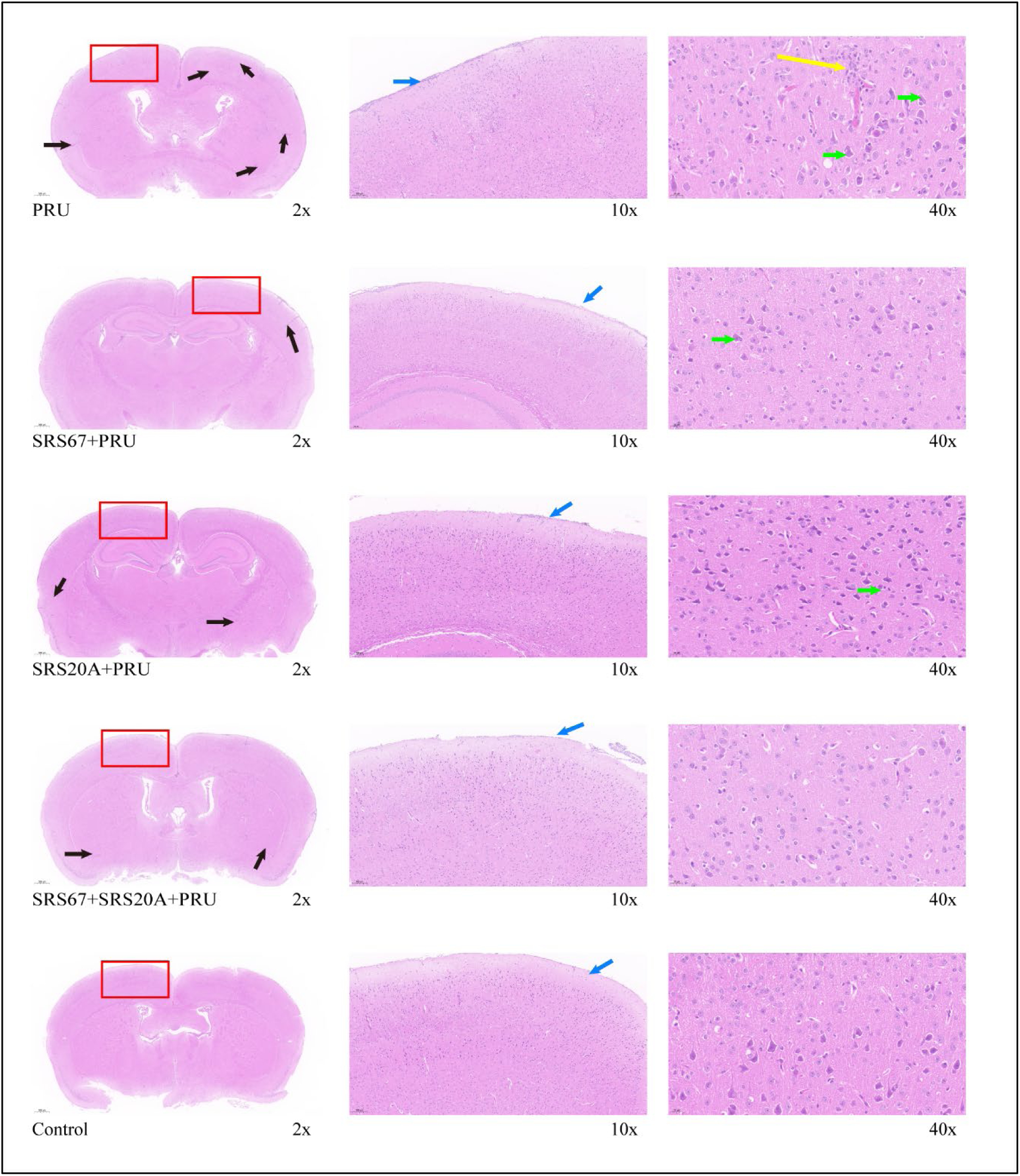
Effect of SRS67 and SRS20A proteins on PRU infected brain tissue.

### Analysis of Oxidative Stress Markers and Cytokine Gene Expression in Immunized Mice

To further explore the molecular mechanisms underlying the immune protection induced by SRS67 and SRS20A, we determined the relative expression levels of antioxidant stress-related genes (SOD, GSH, CAT) and cytokine genes (IFN-γ, TNF-α, IL-18, IL-10, IL-4) in immunized mice from both the PRU (chronic infection) and RH (acute infection) models. The results are presented as follows:

### *T. gondii* RH Acute Infection Model

#### Antioxidant Stress Response

In comparison with the RH-infected group, the relative expression level of the GSH gene was significantly upregulated (*P* < 0.05) in both the SRS67 single-immunization group and SRS20A single-immunization group. Additionally, the relative expression level of the CAT gene was also significantly upregulated (*P* < 0.05) in these groups, while no significant difference in SOD expression was noted (Fig. 6A; P>0.05).

**Fig. 6.**
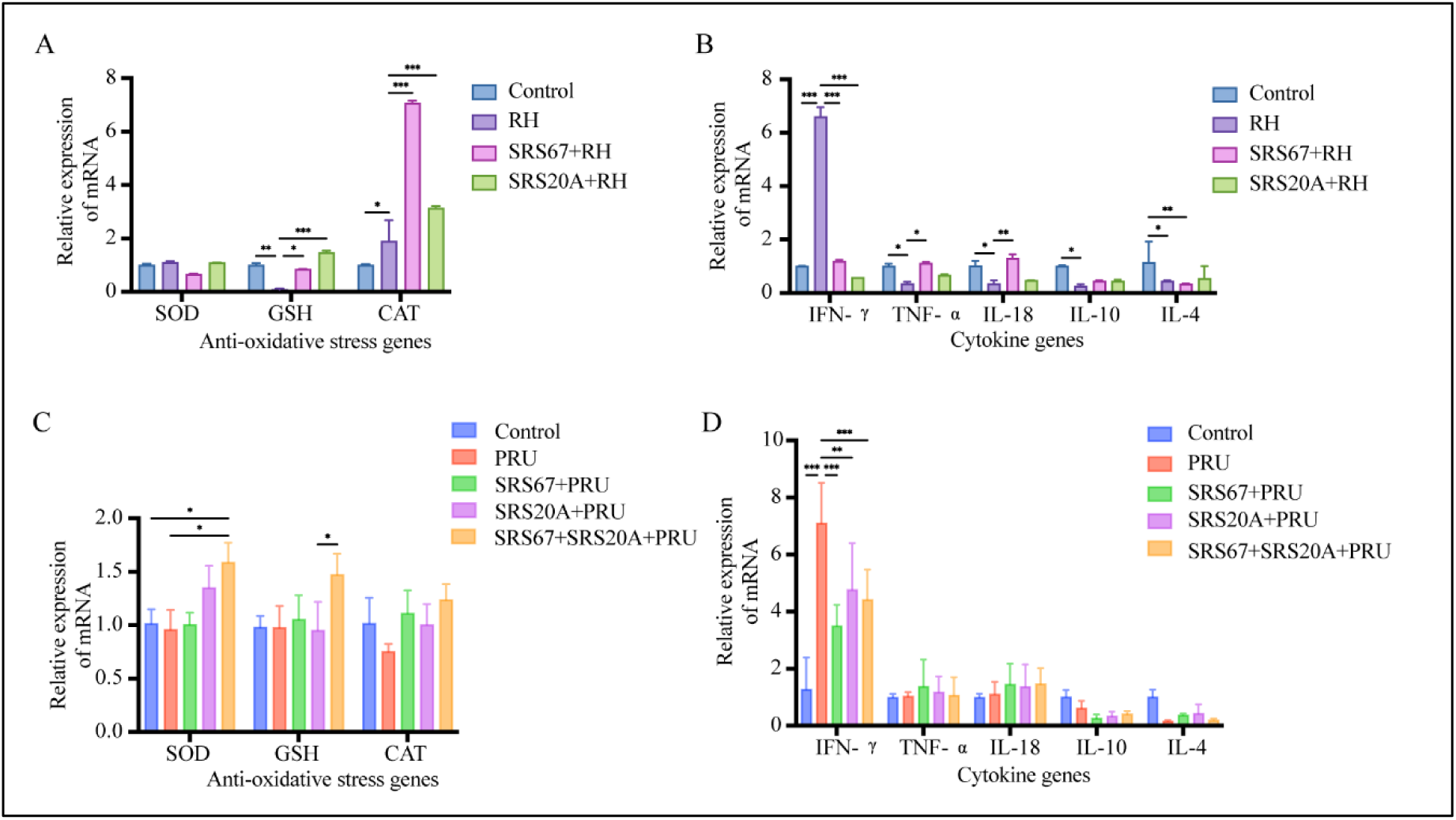
The effects of SRS67 and SRS20A proteins on the oxidative stress-related genes and cytokines in mice infected with both *Toxoplasma gondii* RH strain and PRU strain. A Detection of oxidative stress-related genes in the RH immune protection group. **B** Detection of cytokines in the RH immune protection group. **C** Detection of oxidative stress-related genes in the PRU immune protection group. **D** Detection of cytokines in the Pru immune protection group.

#### Cytokine Response

Compared with the RH-infected group, the relative expression levels of the TNF-α gene (*P*<0.05) and IL-18 gene (*P* < 0.05) were significantly upregulated in the SRS67 singleimmunization group. Additionally, the relative expression level of the IFN-γ gene was significantly downregulated (*P*<0.05) in the SRS67 single-immunization group, SRS20A single-immunization group, and SRS67+SRS20A co-immunization group, while no significant changes were observed in IL-10 and IL-4 expression (Fig. 6B; *P* > 0.05).

### *T. gondii* PRU Chronic Infection Model

#### Antioxidant Stress Response

Compared with the PRU-infected group, the relative expression level of the SOD gene was significantly upregulated (*P*<0.05) in the SRS67+SRS20A co-immunization group. This finding suggests that co-immunization may synergistically enhance the activity of host superoxide dismutase, thereby improving the antioxidant stress capacity. No significant differences in the expression of GSH and CAT were observed between the SRS67 single-immunization group, SRS20A single-immunization group, and the PRU-infected group (Fig. 6C; *P* > 0.05).

#### Cytokine Homeostasis

Relative to the PRU-infected group, the relative expression level of the IFN-γ gene was significantly downregulated (*P* < 0.05) in the SRS67 single-immunization group, SRS20A single-immunization group, and SRS67+SRS20A co-immunization group. In contrast, no significant changes were detected in the expression of IL-10 and IL-4 (Th2-type cytokines) or TNF-α and IL-18 (Th1-type cytokines) (Fig. 6D; *P* > 0.05).

## Discussion

The SRS superfamily comprises the dominant surface antigens of *Toxoplasma gondii,* and their structural and functional properties are essential for host-cell invasion and immune evasion[20]. To elucidate the evolutionary characteristics of two key surface-antigen-encoding loci in the ME49 strain, *TGME49_226860* and *TGME49_285870*, we retrieved the top-scoring homologs from BLAST searches (20 sequences for *TGME49_226860* and 17 for *TGME49_285870*) and reconstructed phylogenetic trees from both nucleotide and amino acid alignments (Fig. 6A–D). Phylogenetic and structural analysis revealed that the homologous sequences of SRS67 and SRS20A of *Toxoplasma gondii* exhibit high conservation, and the nucleotide and amino acid sequences of these homologous regions have over 98% identity, negligible genetic distance (≈ 0), and superimposable three-dimensional folds that each harbour a canonical SAG domain. In cross-species comparisons, *TGME49_226860* orthologs branched successively with *Hammondia hammondi*, *Neospora caninum*, *Cystoisospora suis* and *Besnoitia besnoiti*, whereas *TGME49_285870* formed a well-supported sister group with *N. caninum*; topological incongruence between nucleotide- and protein-based trees was observed. Heat-map profiling of pairwise sequence identity and genetic-distance matrices corroborated these findings (Fig. 6E-F; Fig. S3; Tables S1, S2). Homology modelling indicated that *H. hammondi* and *C. suis* lack the SAG domain and adopt markedly altered conformations, while *N. caninum* retains the domain but displays local structural deviations, underscoring an “intra-species conservation/inter-species divergence” paradigm that presumably fine-tunes host adaptation and functional specialisation across the Apicomplexa. Building on these evolutionary analysis data that *T. gondii* SRS67 and SRS20A are conserved within the species but divergent between species, we have systematically evaluated the immunological profiles of *T. gondii* SRS67 and SRS20A and compared them with established protective antigens SRS34A (Tables S3, S4)[21], SRS47D[16], GRA12[22] and ROP8[23]. Immunological analyses were conducted following the standardized protocols established in our laboratory[19]. Both SRS67 and SRS20A are rich in B-cell, Th-cell and CTL epitopes and exhibit robust immunoreactivity; the immunodominance of SRS20A aligns with previous reports[24,25], whereas SRS67—although less studied—has an *N. caninum* orthologue that elicits *T. gondii*-specific antibodies without cross-reactivity[26,27], highlighting its promise as a broad-spectrum vaccine candidate.

The *T. gondii* type II PRU strain is classically associated with chronic infection and tissue-cyst formation, whereas the type I RH strain is renowned for acute virulence[28]. Quantitative PCR revealed that the transcript abundance of SRS67 in PRU was significantly higher than in RH (P < 0.001), while SRS20A expression did not differ between the two strains. This indicates that SRS20A probably subserves a housekeeping role—e.g., initial attachment and invasion—rather than a stage-specific immunomodulatory function. In contrast, SRS67 was markedly upregulated in bradyzoites versus tachyzoites, implying a contribution to cyst wall biogenesis or maintenance[29–32]. Transcriptomic surveys have shown that >50% of PRU genes are reprogrammed during stage conversion, a process largely driven by host-parasite crosstalk[33,34]. Thus, the robust expression of SRS67 in the PRU strain may reflect an adaptation for long-term cerebral persistence and/or immune evasion. CRISPR-Cas9 knockout of SRS67 in PRU followed by murine neurocyst enumeration is currently underway to test this hypothesis.

IFA disclosed that both SRS67 and SRS20A partially colocalize with GAP45, predominantly localizing to the parasite surface with a small subset distributed in intracytoplasmic puncta (Fig. 2). The canonical view posits that SAG proteins are uniformly GPI-anchored to the plasmalemma, forming a dense surface coat that mediates invasion and shields against host immunity[35]. The predominant membrane signal observed here is consistent with GPI anchorage and a role as adhesion receptors. The additional cytoplasmic pool may arise from the unique apicomplexan secretory route: nascent molecules could be transiently retained in the ER–Golgi intermediate compartment, or alternatively cleaved/exported via exosomes into the parasitophorous vacuole or host cytosol, where they might exert immunoregulatory functions[36]. If substantiated, SRS67/SRS20A would operate as both membrane-anchored receptors and soluble effectors.

In challenge experiments, mice vaccinated with either SRS67 or SRS20A alone, or with the two antigens combined, exhibited 60% survival against lethal RH infection, whereas negative control succumbed (*P* < 0.01). Against the PRU strain, both monovalent and bivalent regimens conferred 100% survival and markedly reduced cerebral cyst burdens, the bivalent group showing the lowest cyst density. Histopathology revealed that combined vaccination significantly attenuated microglial activation and leukocyte infiltration, as microglial activation and leukocyte infiltration are typical manifestations of the cellular immune response, suggesting that protection is mediated not only by antibody but also by a Th1-biased cellular response[37]. Furthermore, the data on oxidative stress and cytokines further unravel the protective mechanisms of SRS67 and SRS20A. In acute RH strain infection, SRS67 upregulates the expression of TNF-α and IL-18 (Th1-type cytokines), which corroborates the hypothesis proposed in the original study that “Th1-biased immunity is involved in mediating protection”. Meanwhile, both proteins downregulate IFN-γ, a process that prevents host tissue damage (e.g., cerebral inflammation) caused by excessive immune activation. In chronic PRU strain infection, the synergistic effect of SOD upregulation induced by co-immunization and IFN-γ downregulation induced by single-antigen immunization not only enhances the host’s antioxidant capacity to counteract oxidative damage resulting from long-term bradyzoite infection but also maintains immune homeostasis to mitigate chronic inflammation. Additionally, the upregulation of GSH in the RH model indicates that both proteins can assist in combating parasitic infection through a non-immune pathway (i.e., antioxidation). This finding expands the scope of the protective effects exerted by SRS family proteins and provides a novel insight for the multi-pathway design of *Toxoplasma gondii* vaccines. Future studies should extend the observation window beyond 30 days to assess durability of memory, map the kinetics of immune responses, and evaluate cross-strain protection. These data will refine antigen formulation and pave the way for a universally effective and long-lasting *anti-Toxoplasma* vaccine.

In conclusion, recombinant SRS67 and SRS20A exhibit high immunogenicity and robust protective efficacy, providing a solid foundation for the development of next-generation subunit vaccines against toxoplasmosis.

## Supporting information

Serum titer determination of SRS67 and SRS20A proteins.

Transcriptional differences of the SRS67 and SRS20A genes between the RH strain and the PRU strain of T. gondii.

Phylogenetic Analysis, Sequence Similarity Analysis, and Domain Prediction of the Homologous Sequences of T. gondii SRS67 and SRS20A Proteins.

The genetic distance table of the SRS67 nucleotide and amino acid genetic evolutionary tree.

The genetic distance table of the SRS20A nucleotide and amino acid genetic evolutionary tree.

Comparison of some proteins antigenicity of T. gondii.

Comparison of the antigenicity of SRS67 and SRS20A proteins with some protozoa.

## Supplementary Information

### Supplementary material 1

**Figure S1:** Serum titer determination of SRS67 and SRS20A proteins. A Serum titer determination results of SRS67 protein. B Serum titer determination results of SRS20A protein.

**Figure S2:** Transcriptional differences of the *SRS67* and *SRS20A* genes between the RH strain and the PRU strain of *T. gondii.* A Transcriptional analysis of *SRS67* in different virus strains. B Transcriptional analysis of *SRS20A* in different virus strains.

**Figure S3:** Phylogenetic Analysis, Sequence Similarity Analysis, and Domain Prediction of the Homologous Sequences of *T. gondii* SRS67 and SRS20A Proteins. A, B Construction of nucleotide and protein phylogenetic trees for the SRS67 homologous sequences. C, D Construction of nucleotide and protein phylogenetic trees for the SRS20A homologous sequences. E, F Heat maps of the nucleotide and protein identity matrices of the homologous sequences of SRS67 (E) and SRS20A (F). In the lower left corner of each heatmap is the amino acid sequence similarity matrix, and in the upper right corner is the nucleotide sequence similarity matrix. G Prediction of protein domains and three-dimensional structures of SRS67 homologous sequences. H Prediction of protein domains and three-dimensional structures of SRS20A homologous sequences.

### Supplementary material 2

**Table S1:** The genetic distance table of the SRS67 nucleotide and amino acid genetic evolutionary tree.

**Table S2:** The genetic distance table of the SRS20A nucleotide and amino acid genetic evolutionary tree.

**Table S3:** Comparison of some proteins antigenicity of *T. gondii*.

**Table S4:** Comparison of the antigenicity of SRS67 and SRS20A proteins with some protozoa.

## Authors’ contributions

XC: Methodology, Validation, Software, Data curation, Visualization, Writing – original draft. TY: Validation, Data curation, Software. WM: Validation. XL: Validation. HL: Software. SN: Validation. MD: Supervision. SZ: Supervision, Writing – review & editing. QL: Funding acquisition, Supervision, Writing–review & editing. DY: Conceptualization, Writing–original draft, Project administration, Supervision, Funding acquisition, Methodology, Resources, Supervision, Writing – review & editing.

## Availability of data and materials

The plasmids and related data are available from the corresponding authors on reasonable request.

## Declarations Ethics ststement

New Zealand White rabbits (2.5 kg) and female BALB/c mice (6–8 weeks old) were obtained from SPF (Beijing) Biotechnology Co., Ltd. All experimental procedures were conducted in strict accordance with the Regulations for the Administration of Laboratory Animals issued by the Ministry of Science and Technology of the People’s Republic of China and were approved by the Animal Ethics Committee of Beijing University of Agriculture (permit no. BUA2023118).

## Consent for publication

All authors reviewed and approved the final manuscript for publication.

