## Supplementary figures and images for "Immunogenicity and Protective Efficacy of *Toxoplasma gondii* SRS67 and SRS20A Proteins in mice"

### Phylogenetic Analysis, Sequence Similarity Analysis, and Domain Prediction of the Homologous Sequences of T. gondii SRS67 and SRS20A Proteins.

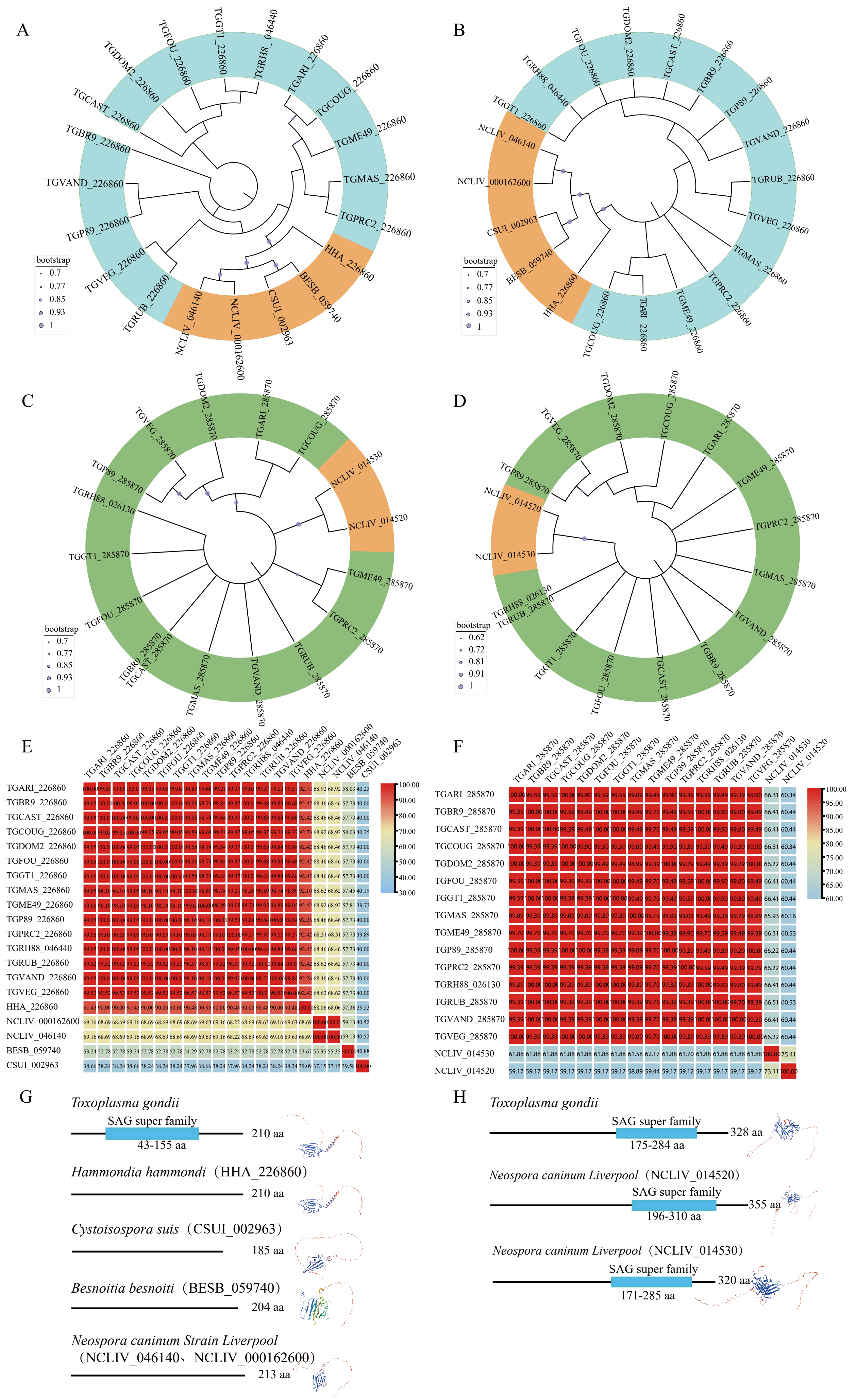

### Serum titer determination of SRS67 and SRS20A proteins.

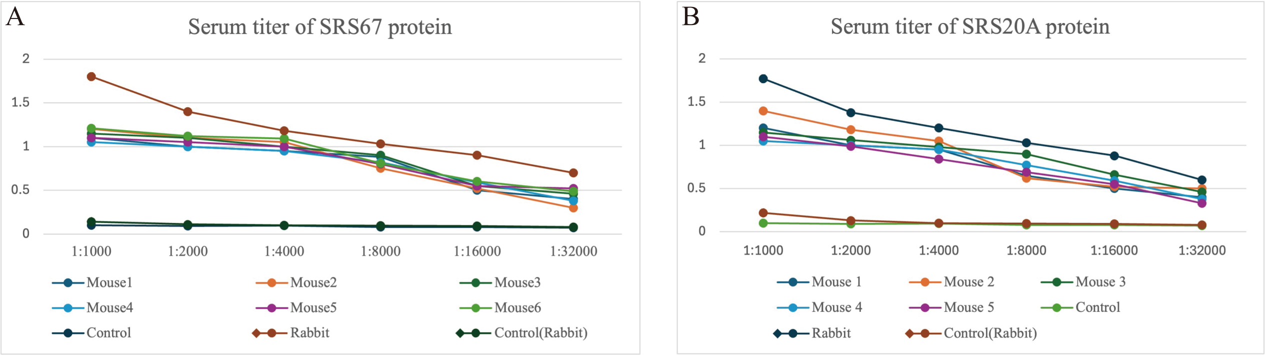

### Transcriptional differences of the SRS67 and SRS20A genes between the RH strain and the PRU strain of T. gondii.

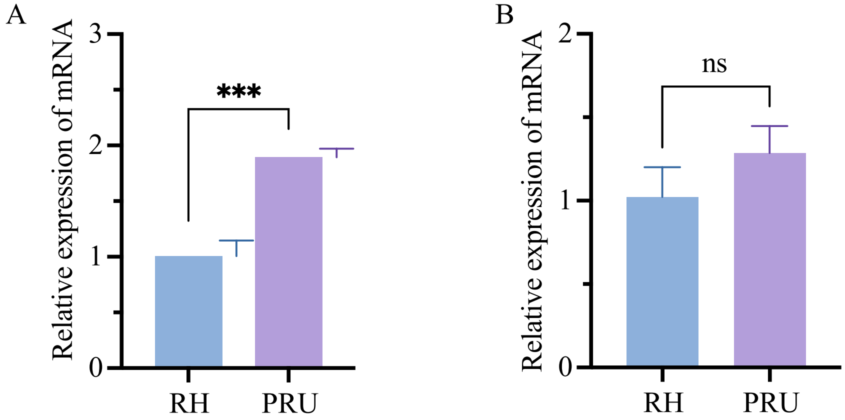
